# From North to South: Transmission Dynamics of H1N1pdm09 Swine Influenza A Viruses in Italy

**DOI:** 10.1101/2024.12.12.628126

**Authors:** Marta Giovanetti, Eleonora Cella, Laura Soliani, Alice Prosperi, Ada Mescoli, Ambra Nucci, Carla della Ventura, Dennis Maletich Junqueira, Nídia S. Trovão, Francesco Branda, Maya Carrera, Davide Lelli, Carlo Rosignoli, Silvia Faccini, Laura Fiorentini, Flavia Guarneri, Gianguglielmo Zehender, Massimo Ciccozzi, Chiara Chiapponi, Ana Moreno

## Abstract

The influenza A H1N1pdm09 virus continues to be a significant pathogen, posing potential risks to both animal and human health due to its zoonotic potential. Italy, which has one of the largest swine populations in Europe, plays a crucial role in monitoring the evolution of influenza viruses in livestock. This study aims to address the existing knowledge gaps regarding the genetic diversity and transmission dynamics of H1N1pdm09 circulating in Italian swine populations. Utilizing whole genome sequencing and dynamic modeling, we conducted a comprehensive analysis of virus samples collected from swine farms across Italy. Our results reveal that multiple independent viral introductions have occurred into the country, with most cases resulting in self-limited infections and limited onward transmission. However, six distinct transmission clusters were identified, suggesting instances of sustained viral spread. These clusters were found across multiple regions of Italy, highlighting the broad geographic distribution of virus lineages. Our findings indicate that while many introductions led to localized containment, certain virus lineages were able to spread within specific regions of Italy. Through a detailed examination of selective pressures, we observed that most viral genes are under strong purifying selection in both swine and human hosts, as reflected by dN/dS ratios well below 1. The hemagglutinin (HA) gene exhibited a notably higher dN/dS ratio in swine (∼0.28) compared to humans (∼0.22), indicating slightly relaxed selection in swine. In contrast, other genes, such as neuraminidase (NA) and non-structural protein (NS), showed similarly strong purifying selection across both hosts. These results reflect a general trend of selective pressures affecting multiple viral components, rather than emphasizing specific genes. Our study emphasizes the importance of ongoing genomic surveillance in detecting viral circulation and mitigating risks to both animal and public health. Italy’s efforts contribute significantly to global influenza monitoring and highlight the importance of a One Health approach that integrates human, animal, and environmental health. These findings provide essential data to inform public health policies and enhance preparedness against future zoonotic influenza outbreaks.

## Introduction

The rise and fall of human history have often been shaped by the appearance and dissemination of infectious diseases (1). Among these, zoonotic diseases, which originate from animals and have the capacity to infect both animals and humans, play a pivotal role. These diseases underscore the profound implications of interspecies interactions, often acting as environments for viral recombination and ushering in new threats with potential epidemic or pandemic dimensions. Within this landscape, the 2009 H1N1 pandemic (H1N1pdm09, clade HA-1A) influenza strains emerged as a significant challenge (2).

At the onset of the 2009 influenza pandemic, the H1N1pdm09 virus was identified as a unique reassortant strain, combining genetic elements from three swine influenza lineages. Notably, key segments were derived from a North American H3N2 swine virus present since the mid-1990s (3). The H1 segment traced back to the classical swine H1N1 lineage circulating since the 1918 pandemic, while the N1 and MP segments were linked to the Eurasian swine lineage that emerged in the late 1970s. The initial human outbreak of the H1N1pdm09 virus is believed to have occurred in Mexico, as indicated by genetic and epidemiological evidence (3). This virus’s ability to cross species barriers is a hallmark of its genetic composition, which includes elements from swine, avian, and human influenza A viruses (4). This characteristic has consistently posed epidemiological challenges. Over time, different H1N1pdm09 strains have emerged, each with unique phenotypic and pathogenic characteristics. Importantly, some of these strains have reached pandemic levels, triggering worldwide health concerns (4).

In European countries, the endemic H1 swine strains circulating since the 1980s, H1avN1 (HA-1C) and H1huN2 (HA-1B), have gradually been joined by new genotypes resulting from various reassortment events, such as novel H1pdm09N2 (HA-1A) and H1avN2 (HA-1C). This has led to increased genetic and antigenic diversity within these H1 lineages (5, 6), contributing to the overall genetic diversity of swine influenza viruses in circulation across European pig herds (7). In Italy, known for its robust pig farming sector, a significant surge in the genetic diversity of swine influenza viruses has been observed over the last ten years (8). This period has seen the emergence of different H1N1 and H1N2 subtypes characterized by diverse gene combinations. Since 2009, Italy has consistently reported the H1N1pdm09 IAV subtype, which accounts for about 10% of the H1N1 isolates identified (9). However, the lack of historical data on the circulation of Influenza A Virus (IAV) in Italian swine populations has made it challenging to trace the origins and transmission of the H1N1pdm09 strain within the country.

Given the importance of pig production in Italy, characterized by a diversified pig population and an extensive herd network, active genomic surveillance is essential. This surveillance is crucial to monitor the evolution of these viruses, particularly regarding reassortment phenomena involving human and avian influenza viruses. Such monitoring will allow for better investigation and understanding of the dynamics of swine influenza and provide crucial insights into global pandemic preparedness and response strategies.

In this study, we employed whole genome sequencing and phylogenetic inference methods to address the existing knowledge gap regarding H1N1pdm09 swine strains circulating in Italy. Our analysis focused on IAV-S strains characterized by the full genome of H1N1pdm09 origin, revealing multiple introductions of the virus, with many resulting in self-limited infections in swine and limited onward transmission. However, we identified a few sustained transmission clusters (n=6) during this period.

Through a comprehensive examination of these strains, including selective pressure analysis (dN/dS), we gained valuable insights into their evolutionary pathways and transmission dynamics within the country. These findings highlight the importance of continuous genomic monitoring as an essential tool for proactive health strategies and emphasize Italy’s significant role in advancing the study of influenza viruses from a One Health perspective.

## Materials and methods

### Ethical approval

Ethical approval was not required for this study as it involved the analysis of residual samples obtained from Influenza A Virus (IAV)-positive specimens (lungs, nasal swabs, or oral fluids). These samples were submitted to the Istituto Zooprofilattico Sperimentale della Lombardia e dell’Emilia Romagna for diagnostic confirmation of pigs with respiratory syndrome between 2009 and 2024. The samples were part of the Italian regional surveillance network, which permits their use for research purposes to accelerate knowledge building and support surveillance and outbreak response efforts.

#### Collection and sequencing of H1N1pdm09 influenza A virus in Italian pigs

Swine influenza sequences were obtained from IAV-positive samples (lungs, nasal swabs, or oral fluids) submitted to the Istituto Zooprofilattico Sperimentale della Lombardia e dell’Emilia Romagna for diagnostic confirmation of pigs with respiratory syndrome from 2009 to 2024. The samples had been previously tested by RT-PCR for the M-gene and subtyped by multiplex RT-PCR (7). Whole genome sequences were obtained from samples with Cts lower than 32, as described previously by Lycett et al. (11), using the SuperScript® III One-Step RT-PCR System with Platinum® Taq High Fidelity (Thermo Fisher Scientific). RT-PCR products were purified with NucleoSpin® Gel and PCR Clean-up (Macherey-Nagel, Carlo Erba, Italy). Sequencing libraries were made with a Nextera-XT DNA Library Prep Kit (Illumina Inc., San Diego, CA, USA) according to the manufacturer’s instructions and sequenced on a MiSeq Instrument (Illumina) using a MiSeq Reagent Nano Kit v2 in a 150-cycle paired-end run. Data were de novo assembled using the CLC Genomics Workbench v.11 (Qiagen, Milan, Italy) and edited with MEGA X (11) and Bioedit 7.2.5 (12). The gene segments were preliminarily analyzed by performing a nucleotide query step in the GenBank database (http://blast.ncbi.nlm.nih.gov/BLAST) to identify closely related sequences. A total of 45 genomes of the H1N1pdm09 lineage were obtained. All samples came from large commercial facilities in the Italian macroregions: Northeast (n=15), Northwest (n=21), Central (n=1), South (n=5), and Insular (n=3). Complete details of all genome sequences obtained in this study are available in **Table S1.**

#### Phylogenetic and phylodynamic analysis of swine H1N1 in Italy and spatial analysis

In addition to the 45 whole-genome sequences generated in this study, background sequences were downloaded from Global Initiative on Sharing All Influenza Data’s (GISAID, https://gisaid.org/publish/) Epiflu database. All available H1N1 pdm09 sequences were downloaded by gene segments. Sequence alignments were constructed for each of the six internal gene segments (PB2, PB1, PA, NP, MP, and NS) and for the two surface antigen segments (HA and NA) using ViralMSA invoking minimap2 (13, 14). Alignments were edited with AliView to remove biological artifacts (15). IQ-TREE2 (16) was used for maximum-likelihood (ML) phylogenetic analysis for each of the eight alignments, employing the general time reversible model of nucleotide substitution and a proportion of invariable sites (+I) as selected by the ModelFinder option. The final datasets for each segment after removing the outlier and low-quality sequences included the following numbers of sequences: HA (n=510, including 45 Italian sequences), NA (n=522, including 45 Italian sequences), PB1 (n=428, including 46 Italian sequences), PB2 (n=459, including 46 Italian sequences), PA (n=481, including 45 Italian sequences), NP (n=464, including 46 Italian sequences), MP (n=554, including 39 Italian sequences) and NS (n=471, including 46 Italian sequences). We performed a joint estimation phylogenetic relationship and the dispersal history for each of the data sets separately using the time-scaled Bayesian approach using MCMC available via the BEAST v1.10.4 (17). A relaxed uncorrelated lognormal (UCLN) molecular clock was used, with a Skygrid population size (18), and a general-time reversible (GTR) model of nucleotide substitution with gamma-distributed rate variation among sites. For computational efficiency the phylodynamic analysis was run using an empirical distribution of 1000 trees (19), allowing the MCMC chain to be run for 100 million iterations (twice, in two separate runs), sampling every 1000. A Bayesian stochastic search variable selection (BSSVS) was employed to improve statistical efficiency for all data sets. All parameters reached convergence, as assessed visually using Tracer v.1.7.1, with all parameters achieving effective sample sizes greater than 200. At least 10% of the chain was removed as burn-in and runs were combined using LogCombiner v1.10.4 and downsampled to generate a final posterior distribution of 1000 trees that was used in subsequent analyses. All parameter values have statistical uncertainty reflected in values of the 95% highest posterior density (HPD) interval. Maximum clade credibility (MCC) trees were summarized using TreeAnnotator v1.10.4, and the trees were visualized using ggtree package in R (20, 21). To identify swine transmission clusters in Italy, we used specific parameters including a bootstrap support threshold of 80% and a mean genetic distance threshold of 1% under the Cluster Picker v. 1.2.5 software (22). Clusters were defined only if they contained at least three swine sequences, and each identified cluster was manually extended until the nearest node was supported by a bootstrap value higher than 80%. This approach ensured robust cluster identification, allowing us to effectively trace swine transmission dynamics.

### Selective pressure analysis

Non-synonymous to synonymous (dN/dS) rate ratios were estimated for each segment with the SLAC method (23) in Hyphy v.2.2.4 (24), focusing on swine samples isolated in Italy. For comparison we also evaluated human samples isolated in this country. The MEME method (25) in DataMonkey (26) was then used to infer the positively selected sites for each dataset (p < 0.05).

## Results

The H1N1pdm09 virus is progressively demonstrating its ability to spread in pig herds both as a result of human-to-pig spread and as a result of pig-to-pig transmission. Indeed, its ability to adapt to the pig population is less than that of the other IAV-S subtypes that have been circulating in pigs for much longer. This results in the detection of lower number of full H1N1pdm09 strains (n=45) than IAV-S strains. However, over the years, there has been an increase in reassortment events between IAV-S and H1N1pdm09 strains where the latter contributes with all or at least one internal genes of pdm09 origin. From 2009 to 2024 we sequenced 120 IAV-S strains with H1A hemagglutinin. Among them 45 were full H1N1pdm09 derived (Genotype P) while 75 were reassorting strains belonging to both H1N1 and H1N2 subtypes with different gene combinations (27, 28). **Figure 1** and Table 1 showed the characteristics of the internal gene cassette (all genes of avian origin, all pdm origin and at least one with different origin) for each H-N combination.

**Figure 1.**
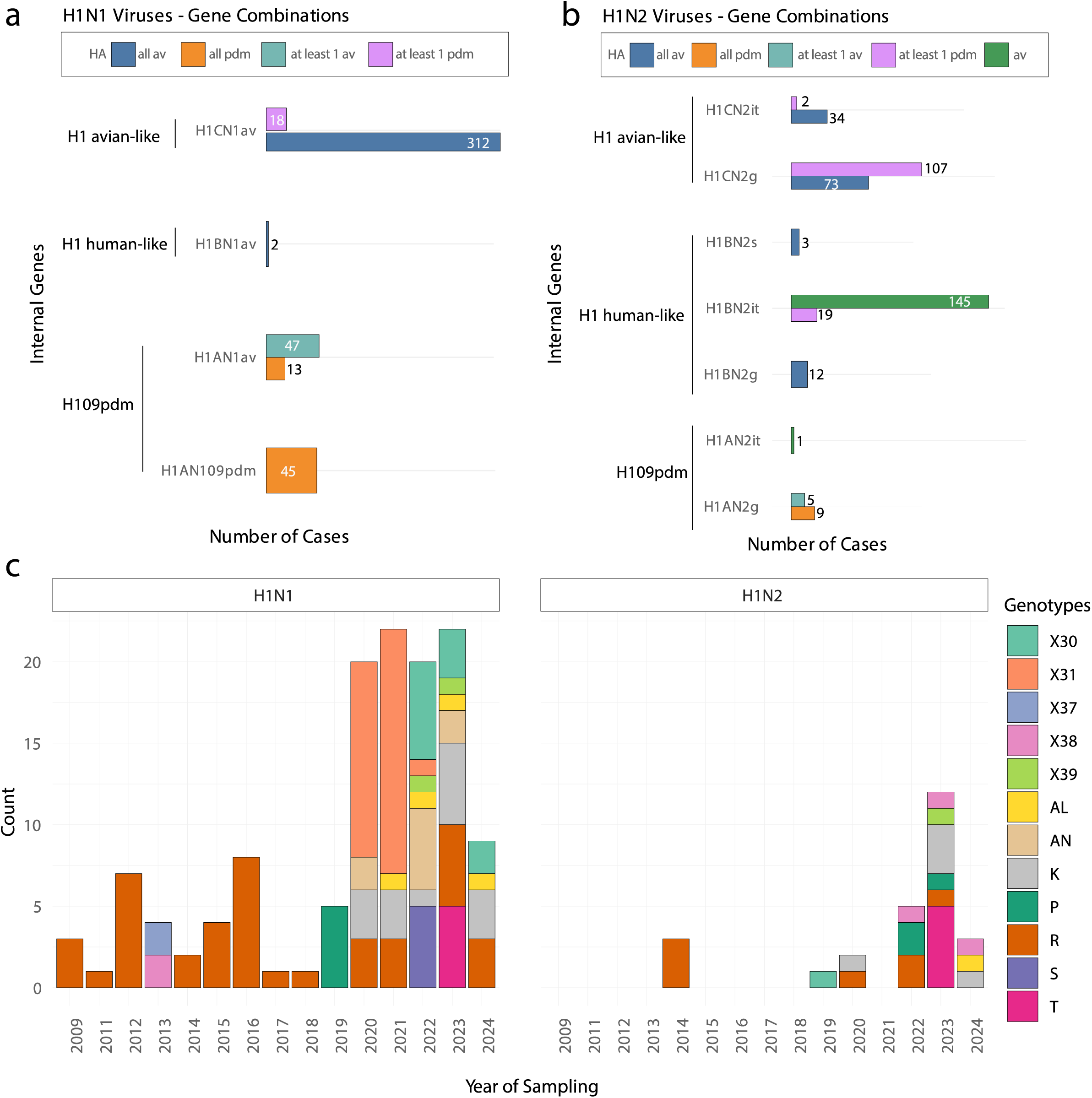
Genetic combinations detected among the H1A origin viruses from 2009 and 2024 in Italy. The origin of each gene segment is described previously (1) and described as pdm (derived from H1N1pdm09) or av (derived from H1N1 av-like) for internal protein coding genes and distinguished into for NA: N1av, N2g or It-N2. Within each subtype, the number of each different genotype (2) detected is described by the genetic sequence HAclade_NA_PB2_PB1_PA_NP_MP_NS.

**Table 1.**
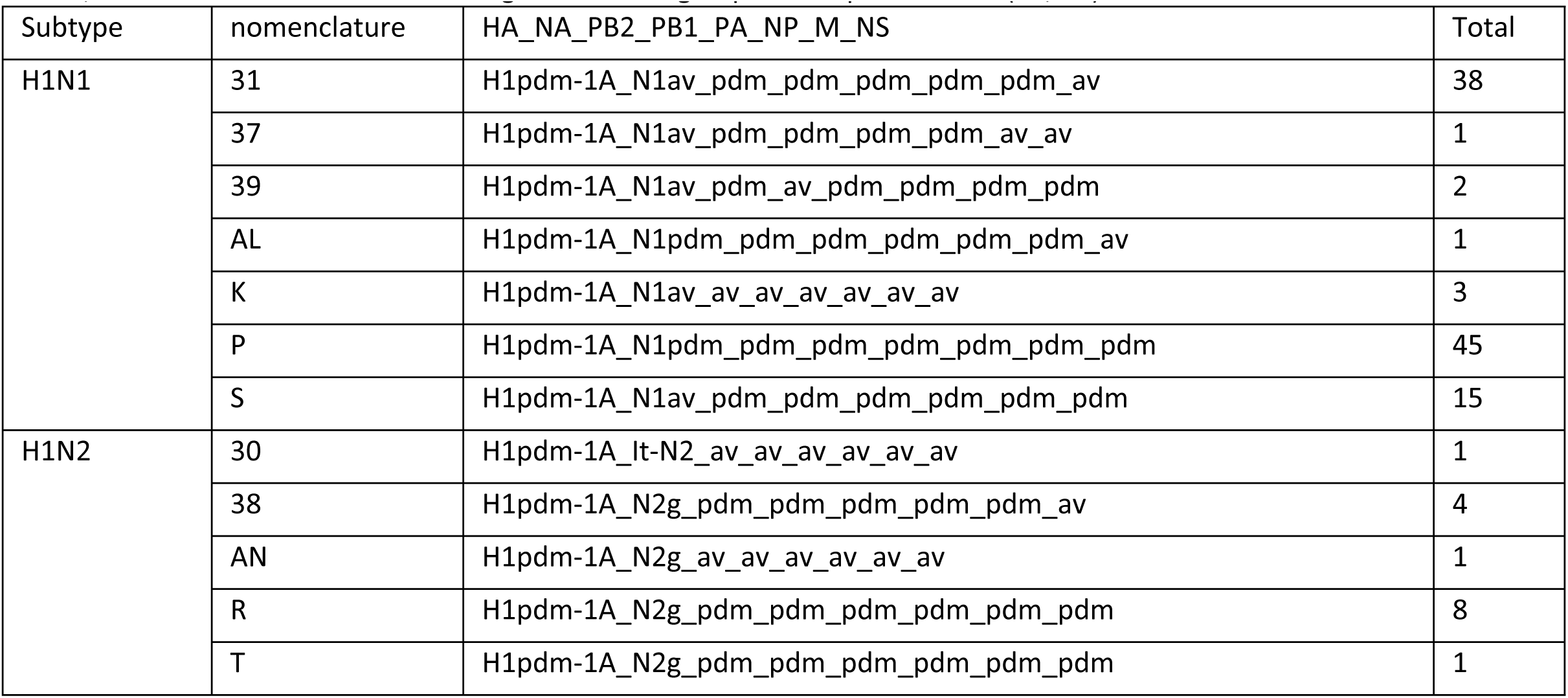
Total number of H1A origin strains sequenced since 2009 with the genotype assigned. Internal gene combination is shown, and nomenclature has been assigned according to previous publications (27, 28).

| Subtype | nomenclature | HA_NA_PB2_PB1_PA_NP_M_NS | Total |
| --- | --- | --- | --- |
| H1N1 | 31 | H1pdm-1A_N1av_pdm_pdm_pdm_pdm_pdm_av | 38 |
|  | 37 | H1pdm-1A_N1av_pdm_pdm_pdm_pdm_av_av | 1 |
|  | 39 | H1pdm-1A_N1av_pdm_av_pdm_pdm_pdm_pdm | 2 |
|  | AL | H1pdm-1A_N1pdm_pdm_pdm_pdm_pdm_pdm_av | 1 |
|  | K | H1pdm-1A_N1av_av_av_av_av_av_av | 3 |
|  | P | H1pdm-1A_N1pdm_pdm_pdm_pdm_pdm_pdm_pdm | 45 |
|  | S | H1pdm-1A_N1av_pdm_pdm_pdm_pdm_pdm_pdm | 15 |
| H1N2 | 30 | H1pdm-1A_It-N2_av_av_av_av_av_av | 1 |
|  | 38 | H1pdm-1A_N2g_pdm_pdm_pdm_pdm_pdm_av | 4 |
|  | AN | H1pdm-1A_N2g_av_av_av_av_av_av | 1 |
|  | R | H1pdm-1A_N2g_pdm_pdm_pdm_pdm_pdm_pdm | 8 |
|  | T | H1pdm-1A_N2g_pdm_pdm_pdm_pdm_pdm_pdm | 1 |

### H1N1 swine phylodynamic reconstruction in Italy

The final dataset of the H1N1 swine genome sequences that we generated and analyzed was from seven of Italy’s twenty regions. The bulk of the sequences originated from the Northwest; Lombardy and Piedmont contributed with 18 and two sequences, respectively. From the Northeast, Veneto provided 11 sequences, and Emilia-Romagna contributed with five. Umbria represented Central Italy with a single sequence. In the South, Campania was responsible for five sequences, and the Insular region, Sicily added three, as shown in **Figure 2a**.

**Figure 2.**
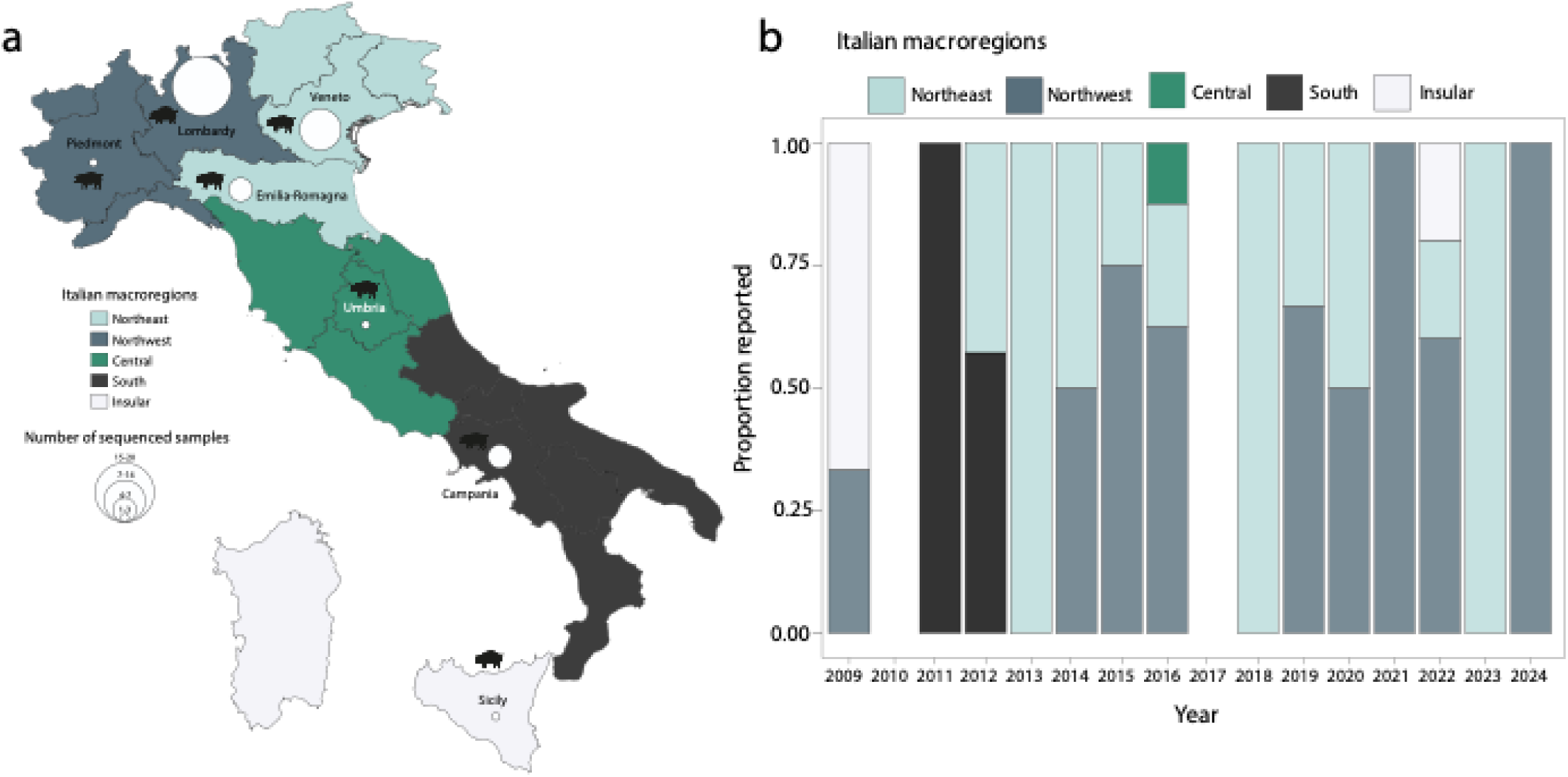
Spatial distribution and genetic diversity of H1N1pdm09 in Italian swine, 2009–2024. a) Distribution of H1N1pdm09 swine genome sequences (n=45) by Italian macro-region. b) Annual count of newly obtained H1N1 swine genome sequences by Italian macro-region. Colors represent different macro-regions: Northeast (light blue), Northwest (blue gray), Central (green), South (black), and Insular (white). Missing years had no genomes.

The temporal distribution of sequences, illustrated in **Figure 2b**, reveals variations in sequencing efforts over the years. The Northeast, particularly Veneto, and the Northwest, especially Lombardy, saw increased sequencing activity after few years, possibly due to a rise in H1N1 cases or enhanced surveillance. During the middle of the study period, the Northwest dominated sequence collection, reflecting its intensive monitoring practices. Meanwhile, Central Italy had minimal representation, and the South and Insular regions, which were more active early in the study period, contributed moderately. This distribution of sequences closely reflects the geographical distribution of pig density throughout Italy, highlighting the need for more consistent and widespread surveillance across all regions to better understand the epidemiology of H1N1 and ensure effective management of the virus in swine populations (29).

To investigate the transmission dynamics of H1N1pdm09 swine influenza strains circulating in Italy, we conducted a comprehensive phylodynamic analysis across the 8 genetic segments, as shown in **Figure 3 and Figure S1**. The analysis identified six distinct clades of H1N1pdm09 circulating within Italian swine populations, marked by recurrent introductions and localized transmission patterns across all genetic segments.

**Figure 3.**
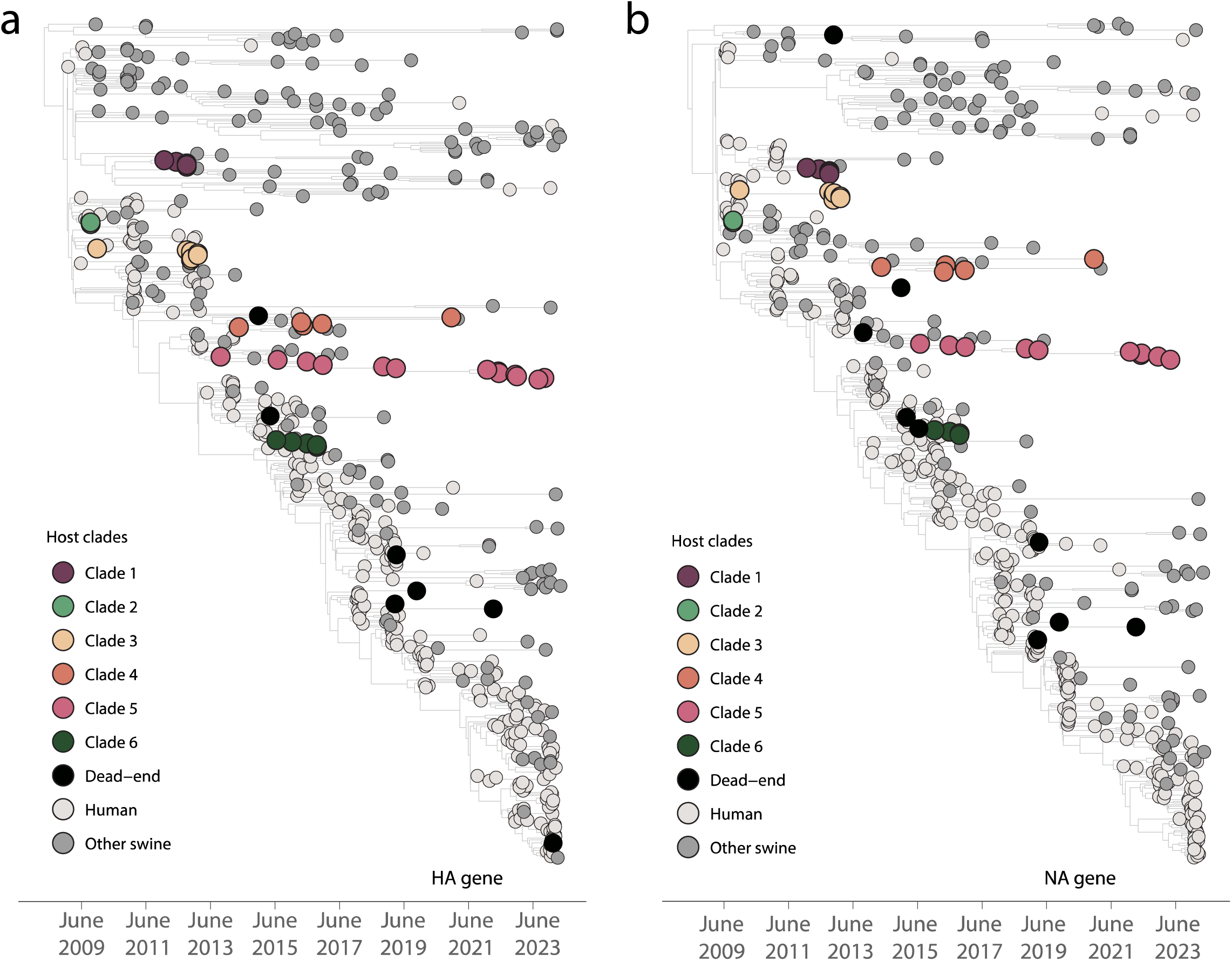
Evolutionary dynamics of Italian H1N1pdm09 HA, NA gene segments. Phylodynamic trees of the HA (a) and NA (b) genes. Tips are color-coded according to clades: Clades 1 to 6 represent monophyletic groups of Italian swine isolates sampled over multiple years, suggesting sustained H1N1pdm09 transmission in Italian swine. Dead-end introductions into Italian swine, which show no evidence of onward transmission, are colored in black. Sequences from humans and swine populations outside Italy are color-coded with light and dark gray, respectively, as indicated in the legend.

Our findings revealed two primary transmissions patterns: self-limited (dead-end) introductions and sustained transmission clusters. Many of the virus introductions, illustrated by singleton swine sequences or small clusters (≤ 2 sequences), represent isolated spillover events where the virus did not establish further transmission. These are considered dead-end introductions, with minimal or no onward spread following the initial infection. This suggests that while frequent human-to-swine transmission events occur mostly from Italy, they often result in contained infections within small, localized groups of swine, following a pattern of sporadic infection that fails to spread widely within the swine population.

We estimated 1 introduction from humans and 3 (1–6) introductions from swine into Italian swine clades.

In contrast, several larger clades, particularly Clades 1, 3, 4, and 5, exhibited sustained transmission over multiple years. These clades are marked by monophyletic groupings of Italian swine sequences, indicating that, once the virus was introduced to swine in certain regions of Italy, it was able to propagate and persist. These sustained transmission clusters highlight the potential for long-term circulation of certain viral lineages, reflecting localized viral adaptation and ongoing transmission within specific swine populations.

Human sequences, interspersed with swine sequences in the phylogenetic trees, confirm that all introductions of the virus into swine originated from human populations, with strong statistical support (pp > 0.8) as observed in the HA and NA genes. As already noted across the gene trees, clustering of swine-origin viruses with sustained swine lineages has been identified across multiple clades, including those for NA, MP, PA, PB2, HA, PB1, NP, and NS. These findings reinforce the importance of monitoring both human and swine populations to capture the full dynamics of interspecies transmission and reduce the risk of future spillovers.

In addition, the phylodynamic analysis reveals distinct regional patterns in the distribution of viral clades across different genetic segments (**Figure S2**). Clade 1 exhibits widespread distribution across the country, with a particularly high concentration of samples from the South (**Figure S2** - black bars), which is present in nearly all gene segments. This clade also circulated in the Northeast (**Figure S2** - light blue), particularly in segments like HA, NS, and PB2, indicating viral transmission among southern and northeastern Italy. Clade 2 was more geographically limited, with most samples coming from the Insular region (**Figure S2** - light grey bars), and it was detectable in the HA and NP gene segments. This clade had little representation elsewhere, suggesting a more confined spread in space and time. Similarly, Clade 3 was sparsely distributed, with its presence mainly restricted to the Central (**Figure S2** - green) and Northwest (**Figure S2** - dark blue) regions, particularly in segments NP and NS, indicating minimal spread beyond these regions. In contrast, Clade 4 showed a broader distribution, with contributions from both the Northeast and Northwest, and some presence in the Central region, particularly for segments like PB1 and PB2. This indicates a wider geographic spread, though its concentration remains higher in northern Italy. Clade 5 stands out as one of the most represented across the country.

Additionally, reassortment events between the HA, NA, and other genome segments of strains circulating in Italian swine were evaluated through phylogenetic inference (**Figure 4, Figures S3 and S4**).

**Figure 4.**
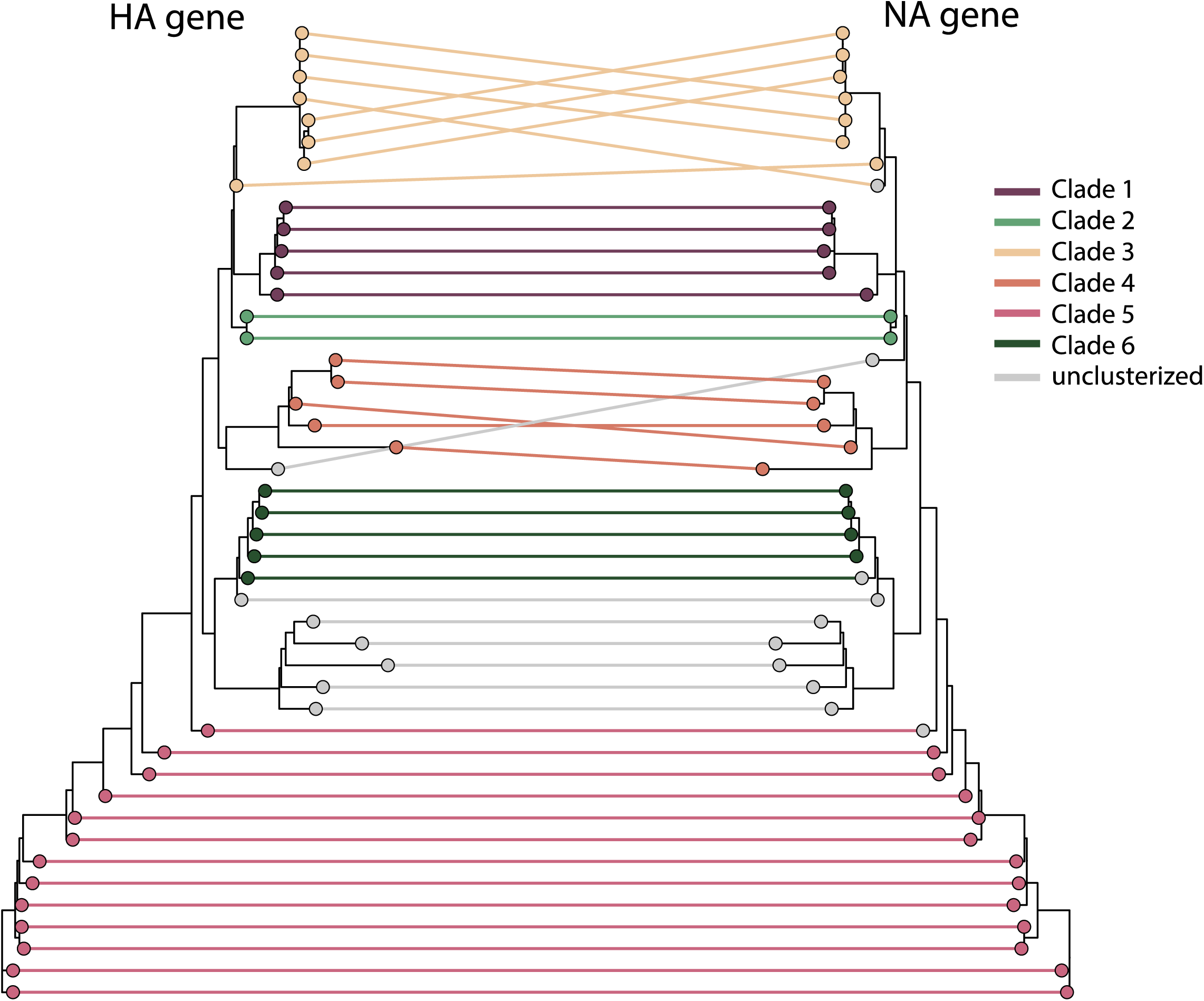
Tanglegram of swine Italian H1N1pdm09 HA, NA strains. Corresponding taxa in the two trees are connected by a line. The tips are colored according to the clade membership. The connecting lines are colored by the HA gene corresponding clade.

Our analysis revealed consistent topologies for HA and NA in Clades 1, 2, 5, and 6. However, we identified two incongruences: one sequence from outside NA Clade 3 grouped within HA Clade 3, and another sequence from outside NA Clades 1, 2, 3, 4, and 6 was found within HA Clade 5. When comparing HA and NA to the other six gene segments, consistent topology and clade assignment were observed only for Clade 2 (**Figure S3**). The greatest number of incongruences were seen in the MP gene comparison, where all clades except Clade 2 showed inconsistencies (**Figure S3**). Most discordances occurred in sequences assigned to a clade in one gene but were unclassified in another but with a similar topology, particularly involving Clades 4, 5, and 6. Mismatched pairings were observed between sequences classified as Clade 5 in one gene and those from Clade 4 in another. This was most notable in comparisons between HA and NP, PB1, and PA, as well as NA and NP, MP, and PA (**Figure S3**). Comparisons of HA and NA with the other six gene segments revealed greater extent of reassortment, while comparing the six segments among themselves showed fewer differences. Notably, significant incongruences were found between MP clades versus the other gene clades (mostly Clade 1 and 4); as well as PB2 clade 4 versus the remaining gene Clade 4 with a discordant topology **(Figure S4**). These findings suggest that reassortment is taking place among H1N1pdm09 viruses circulating in Italian swine, indicating ongoing genetic exchange between different viral strains.

### Evolutionary Dynamics of H1N1pdm09 in Swine Populations Across Italy

In order to gain deeper insights into the evolutionary dynamics of H1N1pdm09 in swine populations across Italy, we applied a thorough phylodynamic analysis. MCC phylogenies were inferred for all HA (N = 6) and NA (N = 6) clades identified in the ML trees (**Figures 5a, 5b**). The obtained phylogenies showed strong consistency with the ML topology, revealing that the majority of persistent introductions into swine originated from common ancestors transmitted from humans between 2009 and 2011.

**Figure 5.**
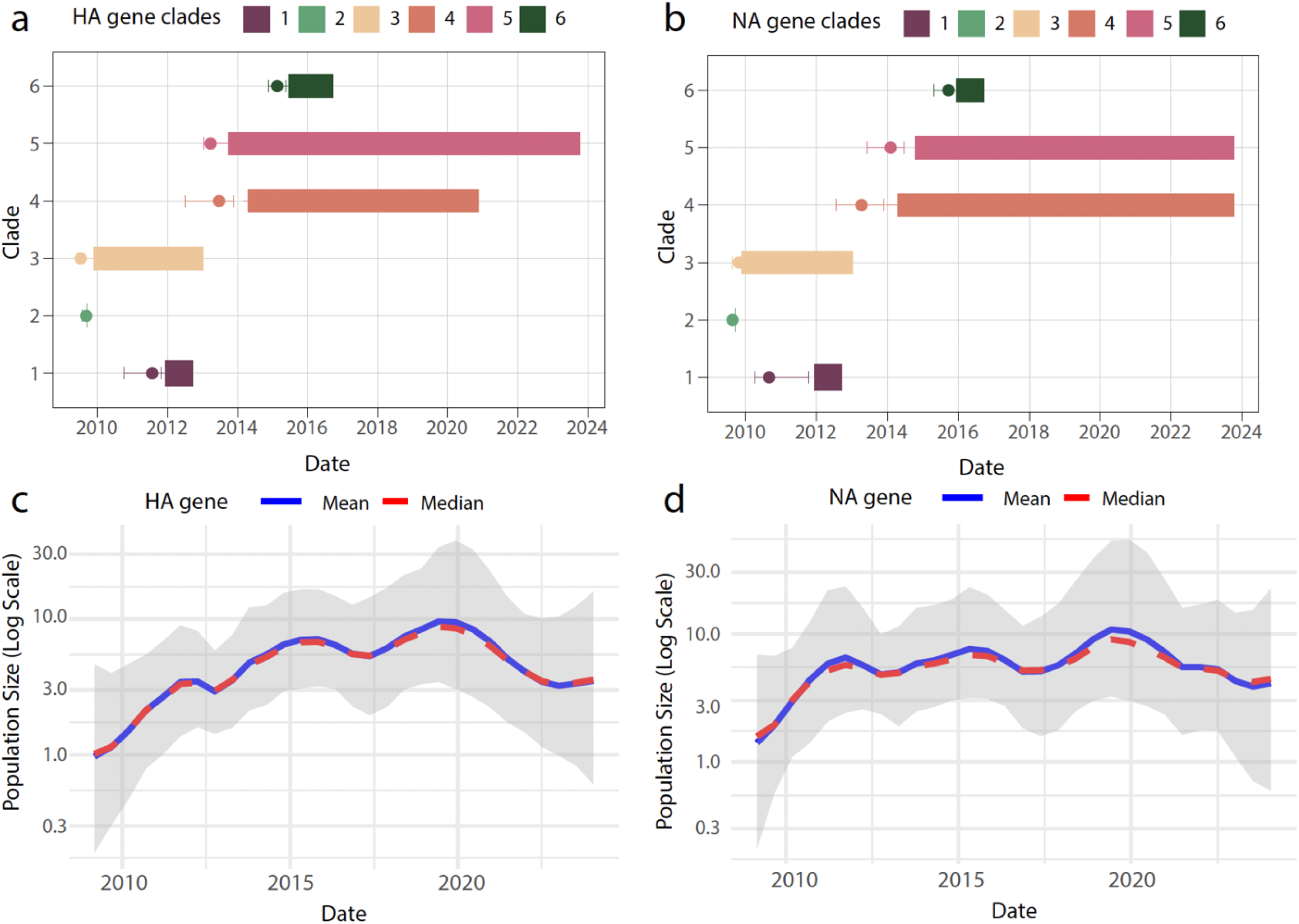
Temporal Evolution and Population Dynamics of H1N1pdm09 HA and NA Gene Clades in Swine Populations. a) Temporal distribution of HA gene clades (Clades 1–6) in H1N1pdm09 swine influenza virus populations in Italy. The x-axis represents the time period from 2010 to 2024, while the y-axis shows the identified clades. Each clade is depicted with colored bars indicating the span of its detection, with markers denoting the median time to the most recent common ancestor (MRCA); b) Temporal distribution of NA gene clades (Clades 1-6), following the same structure as panel (a) c) Bayesian Skyline plot showing the population size dynamics for the HA gene over time, with the effective population size (log scale) on the y-axis and the time on the x-axis. The solid blue line represents the mean estimate, while the red line denotes the median estimate. The shaded area reflects the 95% highest posterior density (HPD) interval; d) Population size dynamics for the NA gene, following the same structure as panel (c).

Bayesian reconstructions suggest that HA Clades 1, 3, 4, 5 likely emerged before 2014 and were detected until 2013, 2013, 2021, and 2023 respectively (**Figure 5a**). Similarly, NA Clades 1, 3, 4, and 5 also emerged before 2014 and persisted in swine populations for several years, Clade 4 and 5 up to 2023 (Figure 5b). Notably, sequences from Clade 5 were found to have circulated in swine for approximately 9 years, highlighting the longevity of certain viral lineages. To further explore the population growth dynamics of H1N1pdm09 circulating in swine, we employed the GMRF Skyride coalescent model (**Figures 5c, 5d**). Analyses of both HA and NA genes revealed an exponential increase in the effective population size (Ne) until approximately 2012, which coincided with the period of predominant human-to-swine transmission (exponential phase). After this phase, the relative genetic diversity stabilized, reflecting a balance between successful swine-to-swine transmissions and viral extinction events (stable phase). Interestingly, a minor decrease in relative genetic diversity was observed from 2020 onwards, suggesting possible reductions in transmission rates or population bottlenecks within Italian swine. To address cryptic transmission, we observed that certain clades, such as Clade 5, circulated for extended periods before being detected, suggesting areas where early detection could be further enhanced. The time between the initial introduction of a clade and its detection varied, reflecting differing rates of cryptic transmission across regions.

### Evolutionary Pressures Shaping H1N1pdm09 within the Italian Swine Populations

We also investigated the selective pressures acting on the H1N1pdm09 virus by comparing the dN/dS ratios (the ratio of nonsynonymous to synonymous substitution rates) across different gene segments in both swine and human hosts (**Figure 6**).

**Figure 6.**
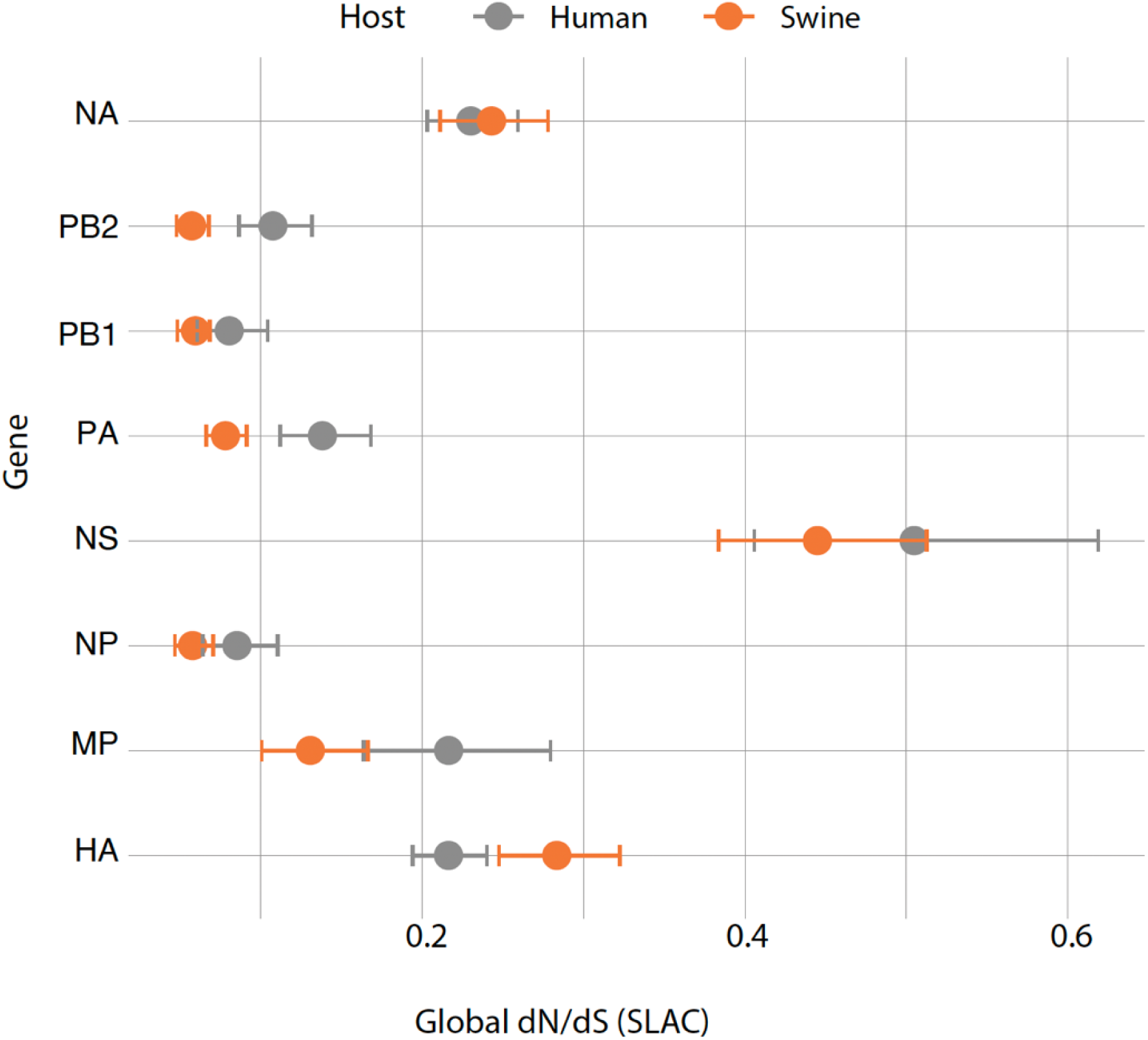
Selective Pressure on H1N1pdm09 Gene Segments in Italian Swine and Human Populations. Global dN/dS ratios (nonsynonymous to synonymous substitution rates) for eight gene segments (NA, PB2, PB1, PA, NS, NP, MP, and HA) of H1N1pdm09 circulating in Italian swine (orange) and human (gray) populations, estimated using the SLAC (Single Likelihood Ancestor Counting) method. The x-axis represents the global dN/dS ratio, and the y-axis lists the gene segments. Circles represent the dN/dS estimates, with horizontal bars indicating the confidence intervals. Ratios below 1 suggest purifying selection.

The dN/dS ratio is an important measure of evolutionary pressure: values below 1 indicate purifying selection (selection against non-synonymous mutations), values around 1 suggest neutral evolution, and values above 1 indicate positive selection (favoring non-synonymous mutations). The analysis reveals all H1N1pdm09 genes are subject to purifying selection in both swine and human hosts, with dN/dS ratios well below 1. These results suggest that these genes are mostly evolving via genetic drift and that non-synonymous mutations are largely deleterious, being selected against in both species. However, some differences between swine and human hosts are observed. The HA gene is the only one exhibiting a higher dN/dS ratio in swine (∼0.28) compared to humans (∼0.22), indicating relaxed selective pressure or adaptation in swine, compared to humans where purifying selection is more pronounced. The overall trend in the dN/dS ratio was similar in both species across all genes, likely due to multiple introductions from humans to swine. These findings highlight that while purifying selection dominates the evolutionary dynamics of the H1N1pdm09 virus in both hosts, the viral attachment protein, HA, may be experiencing weaker selection constraints, particularly in swine. This suggests that swine populations may act as reservoirs for viral evolution, with the potential for genetic changes that may influence transmission and pathogenicity.

## Discussion

The dN/dS ratio is an important measure of evolutionary pressure: values below 1 indicate purifying selection (selection against non-synonymous mutations), values around 1 suggest neutral evolution, and values above 1 indicate positive selection (favoring non-synonymous mutations). Through the analysis of 45 genome sequences from different Italian regions, we gained a deeper understanding of the virus’s evolutionary patterns and its geographical spread within the country. The regional distribution of sequences highlights the uneven nature of surveillance efforts across Italy. Seventy percent of the pig population is concentrated in the northern regions of Lombardy, Veneto, Piedmont, and Emilia-Romagna, where industrial-scale pig farming is predominant (30). This explains the greater number of sequences obtained from these areas, particularly Lombardy, which contributes significantly to national pig production and accounts for a substantial proportion of the sequencing efforts. Conversely, regions with smaller pig populations are naturally expected to yield fewer viral sequences, which may more accurately reflect the real prevalence of circulating strains in these areas.

Addressing this balance in surveillance is key to ensuring a comprehensive understanding of the epidemiology of H1N1pdm09 across the country, as the smaller number of sequences from certain regions may still provide valuable insights into local viral circulation patterns. Strengthening surveillance across all regions, including areas with fewer pigs, could enhance the early detection of emerging strains and support a more proactive approach to influenza monitoring in Italy.

Our phylodynamic analysis revealed a complex pattern of multiple independent introductions of H1N1pdm09 into Italy, leading to both dead-end transmissions and sustained transmission clusters. The detection of six distinct clades, particularly the persistence of Clades 1, 3, 4, and 5, suggests that the virus has been periodically reintroduced, likely from external sources. These reintroductions appear to involve both human and swine populations, establishing swine-specific lineages. This pattern of these introductions underscores the ongoing challenge of managing swine influenza and highlights the need for continuous genomic surveillance to quickly identify and mitigate these events (31). Moreover, our findings highlight important evolutionary dynamics and genetic diversity within the H1N1pdm09 viruses circulating in Italian swine. The consistent topologies observed for the HA and NA genes in Clades 1, 2, 5, and 6 suggest some level of stability in the evolutionary histories of these surface proteins. However, significant incongruences, particularly in sequences outside established NA clades, indicate potential reassortment events. These are further supported by discordances between the HA, NA, and other gene segments, with notable discrepancies in Clades 3 and 5. Such reassortments may give rise to novel viral genotypes with altered pathogenicity or transmissibility, which could impact both swine populations and public health. The incongruences observed in Clade 5 (from the NA comparison) and Clade 3 (from the HA comparison) point to genetic exchanges between different viral strains, potentially resulting in the emergence of new genotypes with modified characteristics. This ongoing genetic exchange, particularly through reassortment, may significantly influence the evolution of swine influenza viruses, potentially leading to changes in pathogenicity, transmissibility, or antigenic properties. Furthermore, the discordances between PA and MP clades, and between PB2 clades and the MP and NS clades, underscore the complexity of the viral evolutionary landscape within swine influenza viruses in Italy.

The selective pressure analysis underscores the importance of host-specific adaptation in shaping the virus’s evolutionary trajectories. All gene segments are under strong purifying selection in both swine and human hosts, reflecting the multiple introductions of strains from humans to swine populations. The use of a vaccine containing H1N1pdm2009 strains in the pig population could also have contributed to this evolutionary trend (32). While most gene segments, including NA, PB2, PB1, and PA, are under strong purifying selection in both swine and human hosts, notable differences were observed in the HA gene, where swine exhibited higher dN/dS ratios. The apparently weaker negative selective pressure in pigs compared to human hosts, indicated by the dN/dS ratio for the HA gene (∼0.28 in pigs versus ∼0.22 in humans), could suggest that genetic changes in this segment are more tolerated in pig populations. Although the difference is small and the ratio remains below 1, this relaxed purifying selection might facilitate the sustained transmission of certain viral clades within these populations.

Our findings also reinforce the importance of a One Health approach (33), integrating human, animal, and environmental health sectors to tackle the multifaceted nature of viral transmission. The evidence of rare evidence of human-to-swine spillovers, coupled with the persistence of swine-adapted lineages, underscores the need for coordinated monitoring of both human and swine populations to fully capture the interspecies dynamics of the virus and to mitigate future zoonotic spillovers. Italy’s significant role in global influenza monitoring is evident, given its large swine population and the ongoing risk of viral reassortment with human or avian influenza strains. By linking veterinary surveillance with public health initiatives, we can better anticipate and respond to viral transmission events that may have pandemic potential. The detection of multiple introductions and sustained viral transmission in Italy’s swine populations further underscores the need for ongoing and widespread surveillance efforts. Increased surveillance is crucial for ensuring a more complete epidemiological understanding. Furthermore, international collaboration is essential to track the global movement of influenza strains and to implement coordinated strategies to manage the spread of zoonotic diseases.

In conclusion, this study provides a comprehensive overview of the transmission dynamics and selective pressures shaping the evolution of H1N1pdm09 in Italian swine populations. The findings highlight the complex interplay between human and swine hosts and emphasize the importance of continuous genomic surveillance, particularly in underrepresented regions. Through collaborative efforts and phylodynamic modeling, we can deepen our understanding of the evolutionary forces driving these changes, assess the risks posed by emerging viral strains, and improve preparedness for future zoonotic outbreaks. By adopting an integrated One Health approach, we can develop more effective and sustainable disease management strategies to mitigate the risks posed by influenza viruses circulating in swine populations.

### Limitation

While our study provides important insights into the transmission dynamics and genetic diversity of H1N1pdm09 in Italian swine populations, several limitations should be noted. First, our sample size, though representative, may not capture the full spectrum of genetic diversity across all Italian swine farms, potentially limiting the generalizability of our findings. Additionally, our analysis is constrained by the inherent limitations of dynamic modeling, which relies on assumptions that may not account for all ecological and epidemiological complexities influencing virus transmission. Finally, while selective pressure analysis offers valuable perspectives on evolutionary dynamics, interpretation should be approached with caution as certain selective signals might be influenced by sampling biases or localized viral adaptations. Addressing these limitations in future studies could help refine our understanding of H1N1pdm09 spread and persistence in swine populations, further enhancing our preparedness against zoonotic influenza risks.

## Supplementary material

**Figure S1.**
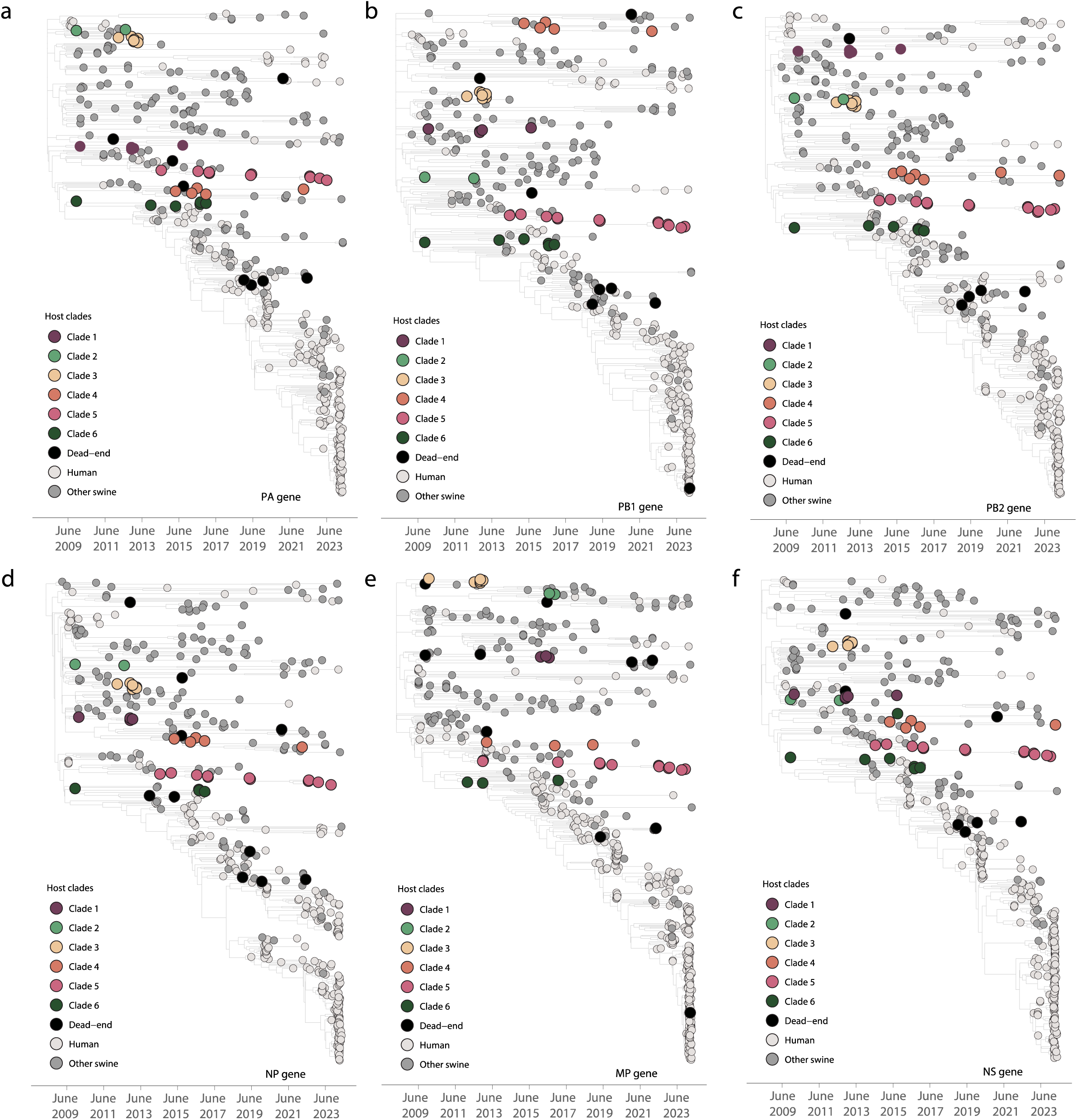
Evolutionary dynamics of Italian H1N1pdm09 PA, PB1, PB2, NP, MP and NS strains. Phylodynamic maximum clade credibility trees of the PA (a), PB1 (b), PB2 (c), NP (d), MP (e), and NS (f) genes. Tips are color-coded according to clades: Clades 1 to 6 represent monophyletic groups of Italian swine isolates sampled over multiple years, indicating introduction and sustained H1N1pdm09 transmission in Italian swine. Dead-end introductions, which show no evidence of onward transmission, are colored in black. Sequences from humans are color-coded in light gray, and sequences from other swine populations are shown in dark gray, as indicated in the legend.

**Figure S2.**
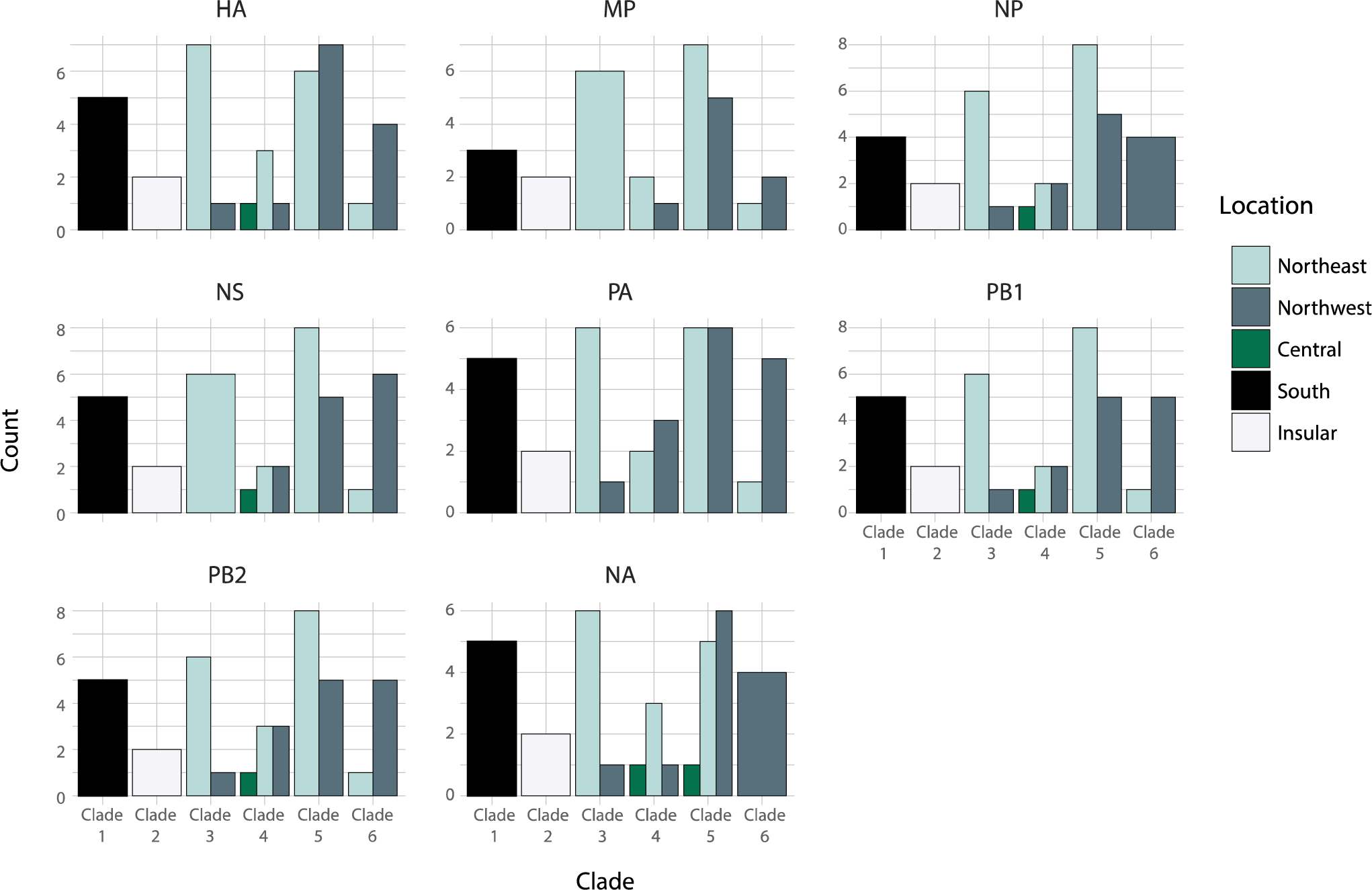
Regional composition of the six viral clades across various genetic segments. The Italian regions are grouped by macroregion and each bar is colored according to the legend on the right.

**Figure S3.**
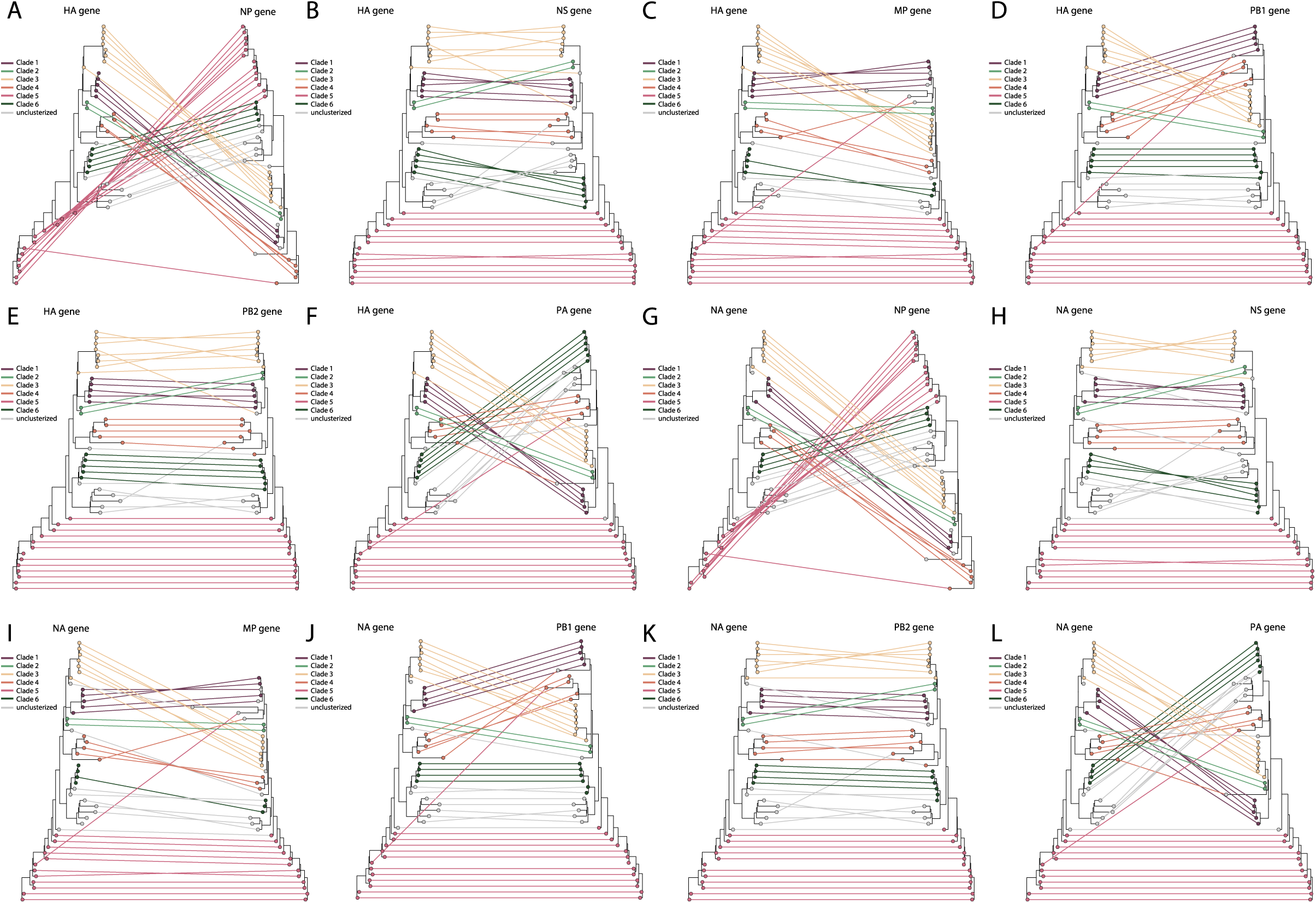
Tanglegrams of swine Italian H1N1pdm09 HA and NA strains with the six internal genes (NS, NP, MP, PA, PB1 and PB2) Corresponding taxa in the two trees are connected by a line. The tips are colored according to the clade membership. The connecting lines are colored by the left-side gene corresponding clade. The legend is on the left-side of the each tanglegram.

**Figure S4.**
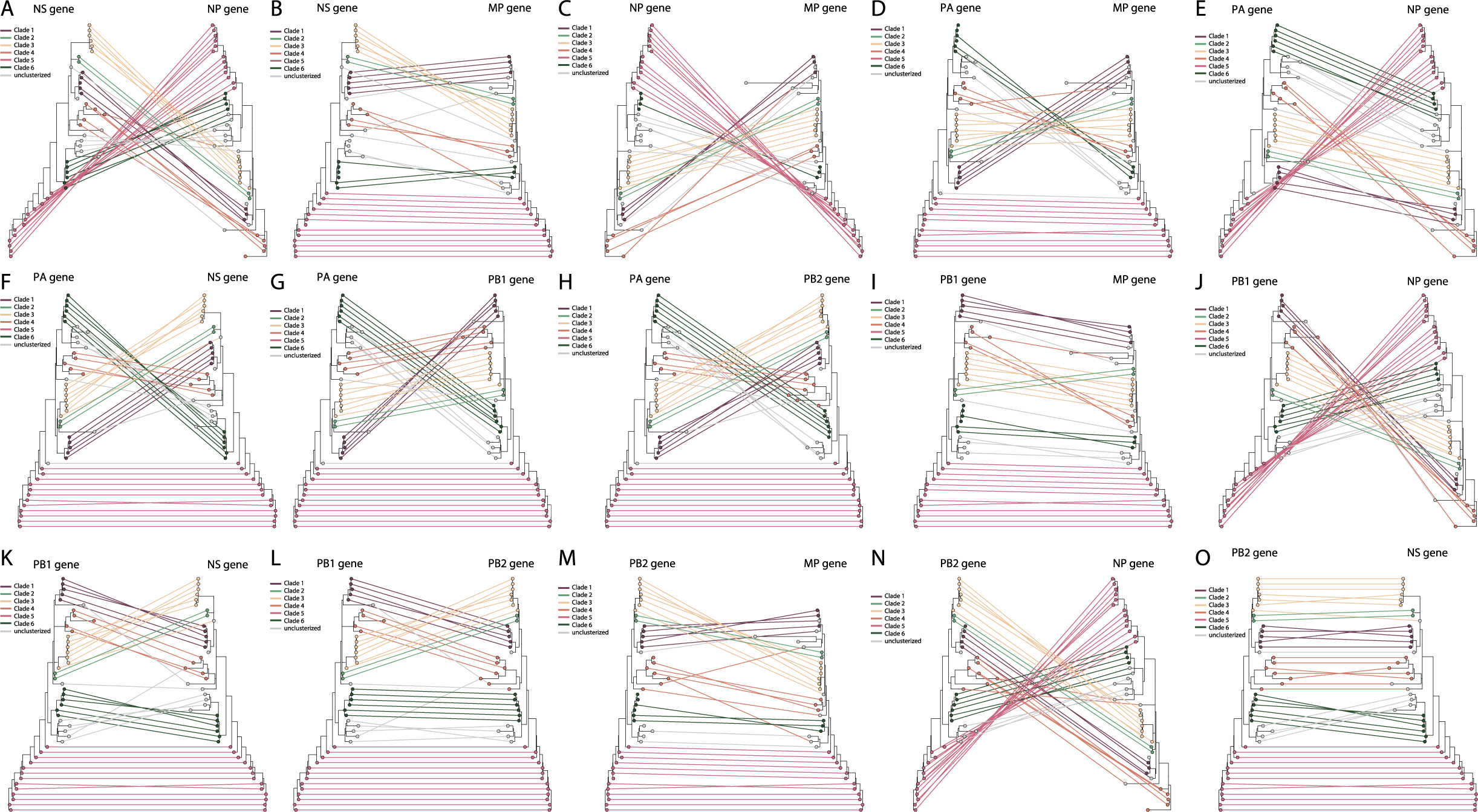
Tanglegrams of swine Italian H1N1pdm09 internal genes (NS, NP, MP, PA, PB1 and PB2) Corresponding taxa in the two trees are connected by a line. The tips are colored according to the clade membership. The connecting lines are colored by the left-side gene corresponding clade. The legend is on the left-side of the each tanglegram.

**Figure S5.**
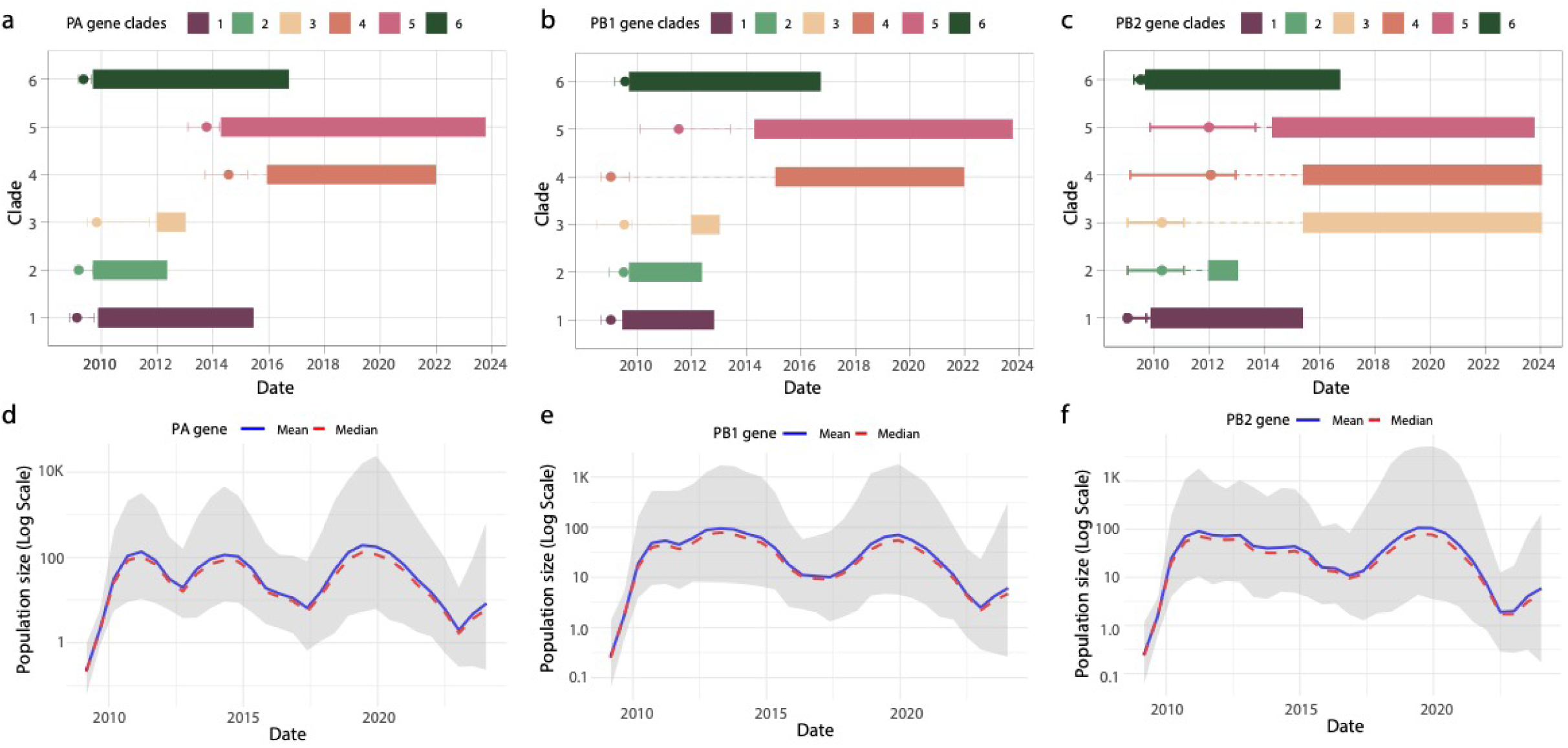
Temporal Evolution and Population Dynamics of H1N1pdm09 PA, PB1 and PB2 Gene Clades in Swine Populations. a) Temporal distribution of HA gene clades (Clades 1–6) in H1N1pdm09 swine influenza virus populations in Italy. The x-axis represents the time period from 2010 to 2024, while the y-axis shows the identified clades. Each clade is depicted with colored bars indicating the span of its detection, with markers denoting the median time to the most recent common ancestor (MRCA); b) Temporal distribution of NA gene clades (Clades 1-6), following the same structure as panel (a), showing the persistence and detection of clades from 2010 to 2024; c) Bayesian Skyline plot showing the population size dynamics for the HA gene over time, with the effective population size (log scale) on the y-axis and the time on the x-axis. The solid blue line represents the mean estimate, while the red line denotes the median estimate. The shaded area reflects the 95% highest posterior density (HPD) interval; d) Population size dynamics for the NA gene, following the same structure as panel (c), showing changes in effective population size (Ne) from 2010 to 2024, with the shaded area representing the 95% HPD interval.

**Figure S6.**
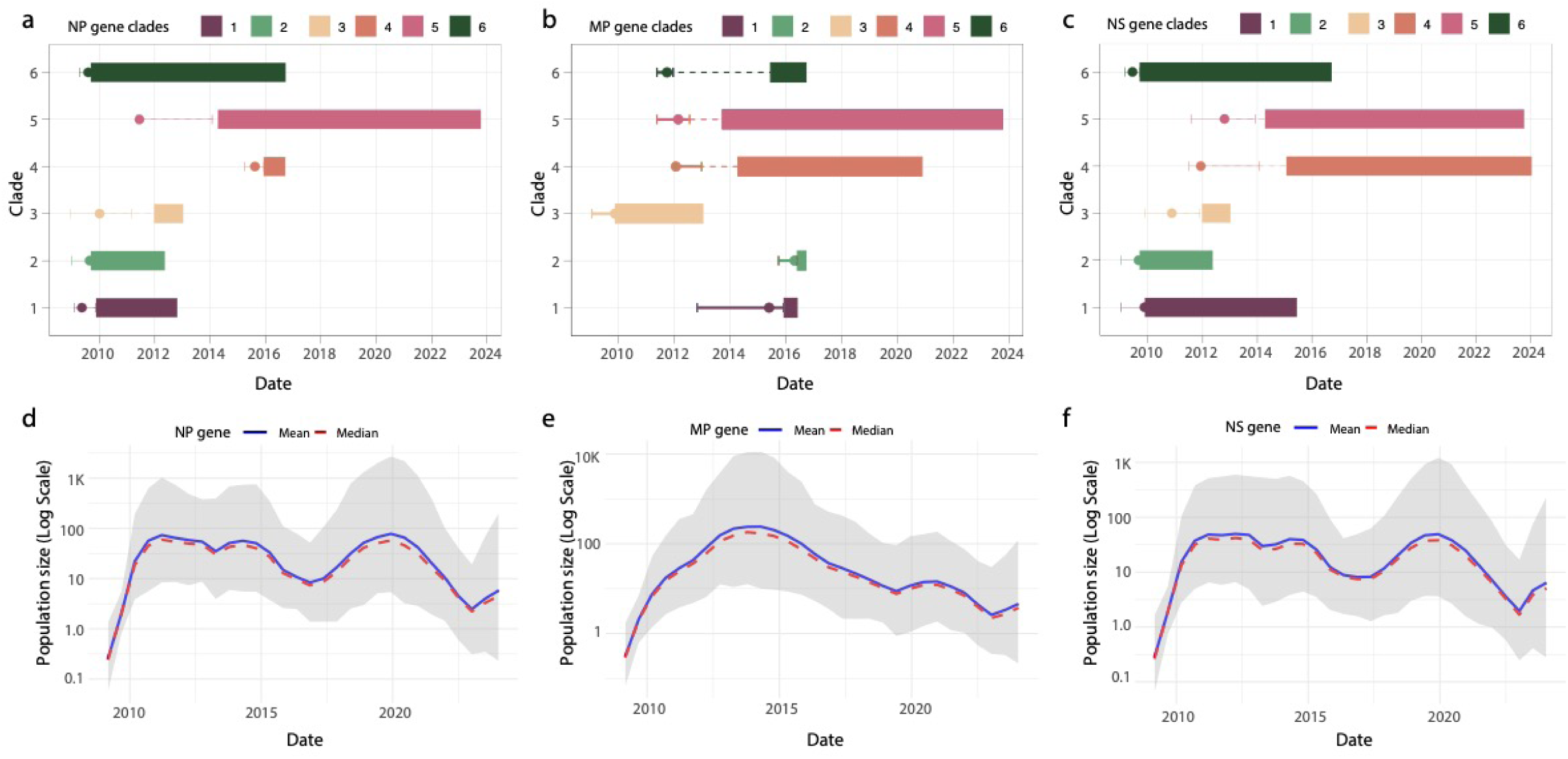
Temporal Evolution and Population Dynamics of H1N1pdm09 NP, MP and NS Gene Clades in Swine Populations. a) Temporal distribution of HA gene clades (Clades 1–6) in H1N1pdm09 swine influenza virus populations in Italy. The x-axis represents the time period from 2010 to 2024, while the y-axis shows the identified clades. Each clade is depicted with colored bars indicating the span of its detection, with markers denoting the median time to the most recent common ancestor (MRCA); b) Temporal distribution of NA gene clades (Clades 1-6), following the same structure as panel (a), showing the persistence and detection of clades from 2010 to 2024; c) Bayesian Skyline plot showing the population size dynamics for the HA gene over time, with the effective population size (log scale) on the y-axis and the time on the x-axis. The solid blue line represents the mean estimate, while the red line denotes the median estimate. The shaded area reflects the 95% highest posterior density (HPD) interval; d) Population size dynamics for the NA gene, following the same structure as panel (c), showing changes in effective population size (Ne) from 2010 to 2024, with the shaded area representing the 95% HPD interval.

## Data Availability

All data associated with the newly generated H1N1 swine Italian genome sequences in this study can be found in **Table S1**.

## Author contributions

Conception and design: M.G., E.C., C.C., and A.M.; Investigations: M.G., E.C., L.S., A.P., A.M., A.N., M.C., D.L., C.R., S.F., L.F., F.G., C.d.A., D.M.J., N.S.T., G.Z., M.C., C.C., A.M., Data Analysis: M.G., E.C., C.d.V., D.M.J., N.S.T., Visualization: M.G., and E.C., Writing – Original: M.G., E.C., C.C., and A.M., Revision: M.G., E.C., L.S., A.P., A.M., A.N., M.C., D.L., C.R., S.F., L.F., F.G., C.d.A., D.M.J., N.S.T., G.Z., M.C., C.C., A.M.

## Acknowledgment

1. M. Giovanetti’s funding is provided by PON “Ricerca e Innovazione” 2014-2020 and by the CRP-ICGEB RESEARCH GRANT 2020 Project CRP/BRA20-03, Contract CRP/20/03. D. Junqueira research’s funding is provided by the State of Rio Grande do Sul (Brazil), through FAPERGS (23/2551-0000879-3). D. Junqueira research’s funding is provided by the State of Rio Grande do Sul (Brazil), through FAPERGS (23/2551-0000879-3). The opinions expressed in this article are those of the authors and do not reflect the views of the National Institutes of Health, the Department of Health and Human Services, or the United States government.

